# Loss of endothelial cell heterogeneity in arteries after obesogenic diet

**DOI:** 10.1101/2023.06.23.546320

**Authors:** Luke S. Dunaway, Melissa A. Luse, Shruthi Nyshadham, Gamze Bulut, Gabriel F. Alencar, Nicholas W. Chavkin, Miriam Cortese-Krott, Karen K. Hirschi, Brant E. Isakson

**Author notes:** to whom correspondence should be addressed University of Virginia School of Medicine, PO Box 801394, Charlottesville, VA 22908 E. these authors contributed equally to this work.

## Abstract

**Background:** It is well recognized that obesity leads to arterial endothelial dysfunction and cardiovascular disease. However, the progression to endothelial dysfunction is not clear. Endothelial cells (ECs) adapt to the unique needs of their resident tissue and respond to systemic metabolic perturbations. We sought to better understand how obesity affects EC phenotypes in different tissues specifically focusing on mitochondrial gene expression.

**Methods:** We performed bulk RNA sequencing (RNA-seq) and single cell RNA-seq (scRNA-seq) on mesenteric and adipose ECs isolated from normal chow (NC) and high fat diet (HFD) fed mice. Differential gene expression, gene ontology pathway, and transcription factor analyses were performed. We further investigated our hypothesis in humans using published human adipose single nuclei RNA-seq (snRNA-seq) data.

**Results:** Bulk RNA-seq revealed higher mitochondrial gene expression in adipose ECs compared to mesenteric ECs in both NC and HFD mice. We then performed scRNA-seq and categorized EC clusters as arterial, capillary, venous, or lymphatic. HFD decreased the number of differentially expressed genes between mesenteric and adipose ECs in all subtypes, but the largest effect was seen in arterial ECs. Further analysis of arterial ECs revealed genes coding for mitochondrial oxidative phosphorylation proteins were enriched in adipose compared to mesentery under NC conditions. In HFD mice, these genes were decreased in adipose ECs becoming similar to mesenteric ECs. Transcription factor analysis revealed C/EBPα and PPARγ, both known to regulate lipid handling and metabolism, had high specificity scores in the NC adipose artery ECs. These findings were recapitulated in snRNA-seq data from human adipose.

**Conclusions:** These data suggest mesenteric and adipose arterial ECs metabolize lipids differently and the transcriptional phenotype of these two vascular beds converge in obesity, in part, due to downregulation of PPARγ and C/EBPα in adipose artery ECs. This work lays the foundation for investigating vascular bed specific adaptations to obesity.

## Introduction

Endothelial dysfunction is prevalent in obesity and is associated with atherosclerosis, myocardial infarction, and peripheral artery disease.^1–5^ However, the etiology and progression to endothelial dysfunction in arterial endothelium is not clear. Additionally, endothelial cells (ECs) adapt to the unique needs of their resident tissue and therefore likely have a heterogenous response to metabolic perturbations, such as obesity.

Recently, there has been much debate over the dependence of ECs on mitochondrial respiration. Early work in this field suggested ECs primarily rely on glycolysis.^6–8^ More recent work has suggested mitochondrial respiration is necessary for fatty acid uptake^9^, angiogenesis^10^, and regulating arterial tone^11^. However, the heterogeneity of EC mitochondrial content between tissues and along the vascular tree remains unstudied.

We investigated EC mitochondrial heterogeneity in the context of a metabolic challenge, high fat diet (HFD). We hypothesized genes involved in mitochondrial function (specifically oxidative phosphorylation) would be differentially expressed between tissues and dysregulated by HFD. In order to investigate changes in endothelial mitochondria in obesity in a tissue specific manner, we performed bulk RNA sequencing (RNA-seq) and single cell RNA-seq (scRNA-seq) on isolated mesenteric and adipose ECs isolated from mice fed normal chow (NC) or HFD. We additionally used publicly available human single nuclei RNA-seq (snRNA-seq) data^12^ to investigate changes in adipose endothelial mitochondrial gene expression in obese human subjects.

## Methods

### Animals

All experiments were approved by the University of Virginia Animal Care and Use Committee and followed the National Institutes of Health guidelines for the care and use of laboratory animals. The mice used in these experiments were a kind gift from Dr. Gary Owens. At 6 weeks old, male ROSA26-eYFP^+/+^ Cdh5-CreER^T2+^ mice were fed a tamoxifen diet (Envigo TD 130856) for 2 weeks to induce YFP expression in endothelial cells. Wet food was provided 3 times a week to ensure adequate consumption. At 8 weeks, the mice were fed either normal chow (NC; 5.8 % fat; Envigo TD 7012) or high fat diet (HFD; 60% fat; Bio-Serve F3282) for 12-13 weeks. Animals were euthanized under CO_2_, and euthanasia was confirmed by cervical dislocation.

### Flow Sorting

Mesentery, epididymal fat pads, and lungs were dissected and placed in ice cold FACS buffer (1% BSA, 3% FBS, 96% Ca^2+^ and Mg^2+^ free PBS) with 1 μg/ml Actinomycin D (ThermoFisher BP606-5). After isolation, samples were moved to a digestion solution (0.2 mg/mlg Liberase TM in PBS (ThermoFisher 50-100-3280), 1 μg/ml Actinomycin D). Samples were minced and incubated at 37 °C for 1 hour or until adequately digested. The samples were further homogenized by passing through a 1 ml pipette tip and 19 G needle. The samples were incubated for 30 min at 37 °C after which they were diluted to 14 ml with cold FACS buffer and placed on ice. The samples were centrifuged at 400 x g for 10 min. The supernatant was collected and centrifuged for 1000 x g for 10 min. Both pellets were recombined and resuspended in RBC lysis buffer (ThermoFisher 00-4333-57) and incubated for a maximum of 5 min. RBC lysis was stopped by addition of 10 ml FACS buffer. The cells were centrifuged at 800 x g for 8 min. The pellets were resuspended and filtered through 70 μm filters. Tissues from 6 mice were combined in a single sample. The single cell suspensions were centrifuged at 800 x g for 8 min and resuspended in a solution of 3% non-acetylated BSA in PBS. Cell viability was determined with Sytox blue (1 μl/ml; ThermoFisher S34857).

Cells were sorted on the Becton Dickinson Influx Cell Sorter with the assistance of UVA Flow Cytometry Core staff. Cells were gated as shown in **Supplementary Figure 1**. Lung from a Cre^+^ mouse and mesentery from a Cre^-^ mouse were used as positive (YFP only) and negative (no color) controls respectively. This experiment was repeated twice to provide two independent replicates.

### Bulk RNA Sequencing

ECs were isolated from mesentery and adipose tissue as described above. RNA was isolated using RNeasy MinElute Cleanup Kit (Qiagen). PE150 reads were generated with the Illumina NovoSeq platform. Adapters were trimmed by with the BBDuk function of BBMap (v38.57). The paired-end reads were mapped to the mm39 reference genome using STAR (v2.7.9a). Gene counts were generated with FeatureCounts in the Subread package. DESeq2 (v. 1.34.0) was used to identify differentially expressed genes (P_adj_ < 0.05). Ensemble IDs were converted to gene symbols using the org.Mm.eg.db package. Raw data and normalized counts have been deposited in NCBI’s Gene Expression Omnibus and are accessible through the GEO accession number GSE235192 (https://www.ncbi.nlm.nih.gov/geo/query/acc.cgi?acc=GSE235192).

### Single Cell RNA Sequencing and Data Analysis

Endothelial cells were isolated from mesentery and adipose tissue as described above. The generation of single cell indexed libraries was performed by the School of Medicine Genome Analysis and Technology Core, RRID:SCR_018883, using the 10X Genomics chromium controller platform and the Chromium Single Cell 3′ Library & Gel Bead Kit v3.1 reagent. Briefly, around 5,000 cells were targeted per sample and loaded onto each well of a Chromium Single Cell G Chip to generate single cell emulsions primed for reverse transcription. After breaking the emulsion, the single cell specific barcoded DNAs were subjected to cDNA amplification and QC on the Agilent 4200 TapeStation Instrument, using the Agilent D5000 kit. Each of the four samples’ cDNA was used to prepare indexed libraries that were pooled prior to sequencing. A QC run was performed on the Illumina Miseq using the nano 300Cycle kit (1.4 Million reads/run), to estimate the number of targeted cells per sample using the Cellranger 3.0.2 function. The cell estimate enabled the core to re-balance the pooled sample prior to deep sequencing onto the NextSeq 2000, using the P3-100 cycle kit. After run completion, the Binary base call (bcl) files were converted to fastq format using the Illumina bcl2fastq2 software raw reads in fastq files were mapped to the mm10 reference murine genome and assigned to individual cells by CellRanger 5.0.0, and data transferred to the Bioinformatics team for further analysis.

Data from two separate experiments were analyzed in RStudio (2022.07.1) with the Seurat package (4.3.0). Sequencing yielded 17,164 cells with 54,100 features. In order to ensure high quality data, cells were excluded if they contained less than 200 genes, more than 5000 genes, if their transcriptome was more than 5% mitochondrial, and more than 5% hemoglobin beta. Data was combined using SCTransform, normalized, and 3000 variable features were chosen. UMAPs were generated using 20 principal components. Clusters were generated using a resolution of 1. Non-endothelial cells were excluded based on low expression of Pecam1 and Cdh5 as well as high expression of non-endothelial markers (Col1a1, Acta2, Cd3g, Ptprc, Ccr5, Adipoq).

UMAPs of individual endothelial populations were generated by subsetting endothelial populations, using the subset() function in Seurat. New 3000 variable features were selected in the subpopulation. The number of principle components and resolution were chosen appropriate for the population. (arterial ECs: pca = 15, resolution = 0.5; capillary ECs: pca = 20, resolution = 0.2; vein ECs: pca = 0.2, resolution = 0.2; lymphatic ECs: pca = 15, resolution = 0.3). Differential gene expression was determined using FindMarkers(). Genes were considered differentially expressed if the adjusted p-value < 0.05 and avg_log2FC > 1 or < −1. The percent of genes differentially expressed was calculated as the number of genes differentially expressed divided by total number of features expressed in that subset.

Gene ontology analysis was performed with enrichR (3.1). Genes that were found to be differentially expressed between tissues in NC but not HFD were analyzed. The top five biological processes as raked by lowest adjusted p-value are reported. Raw data and processed data have been deposited in NCBI’s Gene Expression Omnibus and are accessible through the GEO accession number (https://www.ncbi.nlm.nih.gov/geo/query/acc.cgi?acc=GSE235192).

### Processing of Previously Published Human Data

We utilized a recently published scRNA-seq and snRNA-seq dataset from human adipose tissue.^12^ Arteries endothelial cells were subsetted, using subset() function, from previously published data using clusters defined by the original authors and confirmed by our arterial endothelial markers. Only arterial endothelial cells from visceral adipose tissue were used for analysis for more appropriate comparison with epididymal adipose ECs. This resulted in a data set only containing snRNA-seq data of 1839 ECs (307 male and 1532 female) from 10 subjects (3 male and 7 female). A module score of all mitochondrial genes were generated using AddModuleScore().

### SCENIC

The Seurat object was converted to an expression matrix format compatible with the single-cell regulatory network inference and clustering (SCENIC; v1.3.1) computational pipeline, with genes represented in rows and cells in columns. SCENIC running on R v4.1.1 was then used to perform gene regulatory network analysis based on the standard vignette.^13^ The analysis was run on a multi-core machine, supported by UVA research computing, to parallelize processing. For both mouse and human cell analysis, seven-species databases that scored motifs in the 500bp region upstream of the transcriptional start site (TSS) and 20kb region around the TSS were used. Genes not present in these databases were filtered out from the expression matrix, along with genes not detected in at least 1% of cells and genes with a total expression across all samples of less than 6 UMI counts. As specified in the SCENIC vignette, the seed 123 was used in all runs to reproduce results with minimal randomness. Results from the SCENIC pipeline included cell-specific regulon activity scores, which predicted the activity of every analyzed transcription factor and its target genes in each cell. These activity scores were aggregated by known cell type and used to calculate a Regulon Specificity Score (RSS) for each regulon, indicating the specificity of the regulon’s activity for each cell type.^14^

### En Face

Third order mesenteric and epidimyl fat pad arteries (100-200µm in diameter) were collected and stripped of adipose and connective tissue. Arteries were subsequently fixed in 4% paraformaldehyde on ice for 10 minutes. For *en face* preparation, arteries were cut longitudinally with microdissection scissors and pinned open on polymerized Sylgard 184 (Electron Microscopy Sciences) using tungsten wire (0.0005”, ElectronTubeStore). Vessels were then permeabilized with 0.5% TritonX100 in PBS for 30 minutes at room temperature and blocked in 1% bovine serum albumin, Fraction V (BSA Sigma) in 0.5% TritonX100/PBS. Primary antibody staining was performed overnight at 4°C in 0.1% BSA in 0.5% TritonX100/PBS. Cytochrome C (Abcam, ab90529, 1:100) primary antibody was used. Samples were incubated with secondary antibodies at 1:500 for 1-2 hours at room temperature. Nuclei were stained using DAPI (Invitrogen, D1306, final concentration 0.1mg/mL) in addition to mounting with Prolong Gold Antifade Mountant (Invitrogen, P36930). Images were collected using an Olympus FV3000 with a 60X oil emersion lens and post processing was completed using FIJI.

### Statistics

Cell distributions and percent differentially expressed genes were analyzed with Chi-squared test and Holms correction where indicated in either RStudio or GraphPad Prism (v9.4.0).

### Data availability

Raw data and processed data generate by bulk RNA-seq and scRNA-seq have been deposited in NCBI’s Gene Expression Omnibus and are accessible through the GEO accession number GSE235192 (https://www.ncbi.nlm.nih.gov/geo/query/acc.cgi?acc=GSE235192).

## Results

*Adipose ECs have higher oxidative phosphorylation gene expression by Bulk RNA-seq* To investigate how mitochondrial gene expression varies in ECs across tissues during obesity, we performed bulk RNA-seq on isolated ECs from the mesentery (Mes) and the epidiymal adipose tissue (Epi) of ROSA26-eYFP Cdh5-CreER^T2+^ mice fed either a NC or HFD for 12 weeks, as described in the methods (Figure 1A). We found adipose ECs had significantly higher expression of genes involved in mitochondrial respiration compared to Mes ECs regardless of diet (Figures 1B **– 1D** and **Supplemental Table 1**).

**Figure 1:**
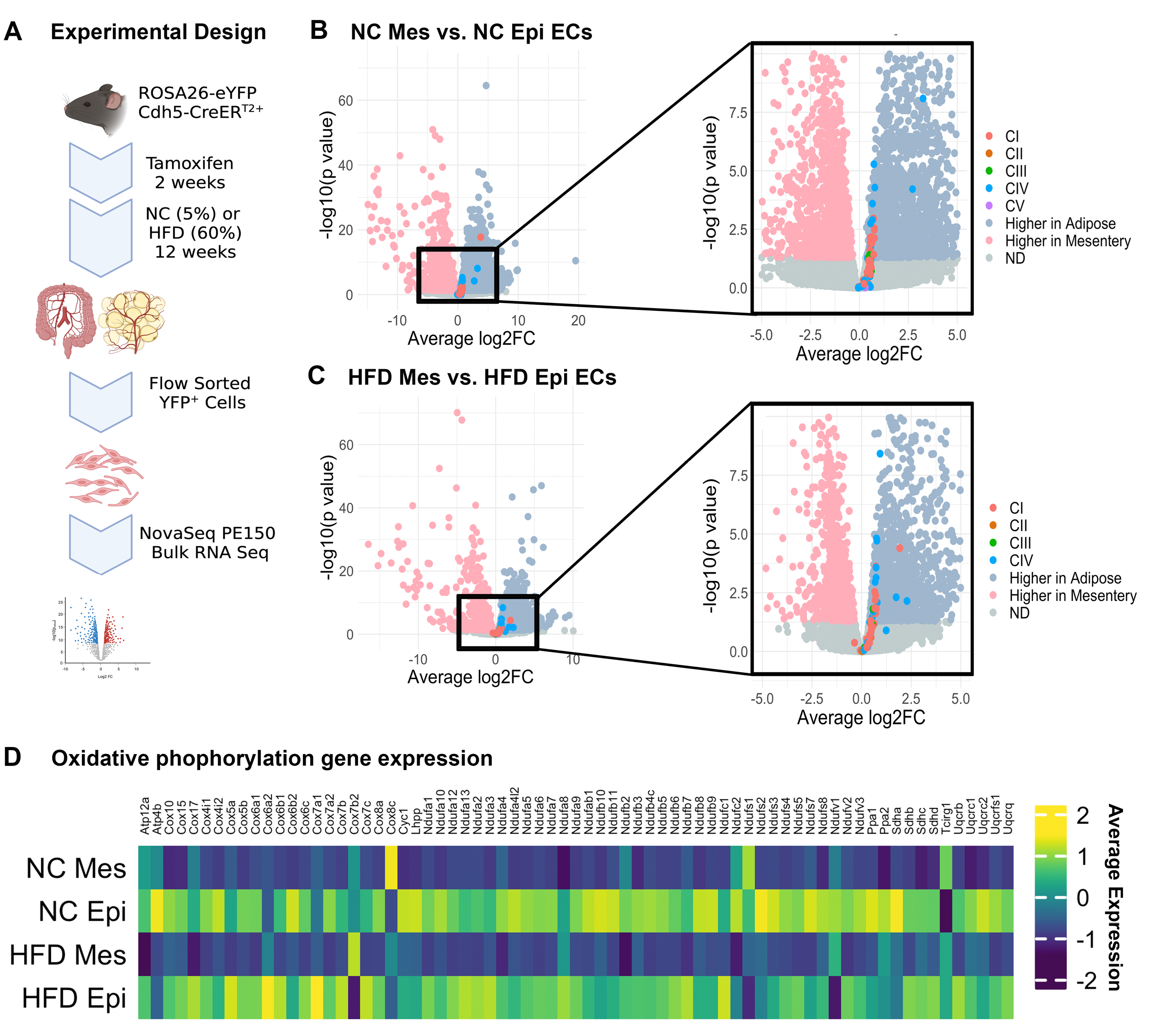
Homogenous populations of adipose ECs have higher oxidative phosphorylation gene expression compared to mesenteric ECs by bulk RNAseq. Experimental design (A) showing the workflow for the isolation of ECs utilizing an inducible YFP reporter mouse under the control of the Cdh5-Cre promoter. Volcano plot (B) showing the differential expression between normal chow (NC) mesenteric (Mes) ECs and NC epididymal (Epi) fat pads. (C) Volcano plots comparing gene expression profiles between high fat diet (HFD) Mes and Epi ECs. (B/C) Insets specifically highlighting an increase in genes associated with oxidative phosphorylation. (D) Heatmap showing several oxidative phosphorylation genes and their expression signature (Color). Bulk RNAseq shows EC expression of oxidative phosphorylation genes is dictated by tissue type. N=3 mice.

### scRNAseq on ECs reveals diet induced transcriptomic convergence

To understand how HFD affects mitochondrial gene expression at a single cell resolution, we performed scRNA-seq on isolated ECs from male ROSA26-eYFP Cdh5-CreER^T2+^ mice fed either a NC or HFD for 12 weeks, as described in the methods (Figure 2A). We identified 16 clusters. Contaminating cell types were identified by low expression of endothelial markers (Cdh5 and Pecam1) and high expression of non-endothelial markers (Col1a1, Acta2, Cd3g, Ptprc, Ccr5, or Adipoq) (Figures 2B and 2C). Contaminating clusters were excluded from further analysis resulting in a dataset of 14,173 ECs. ECs were then further classified based on their expression of genes known to be enriched in arterial (Efnb2, Dll4, Gja5), capillary (Apq7, Rgcc, Gpihbp1), venous (Vcam1, Vwf, Plvap) and lymphatic (Prox1, Lyve1, Flt4) endothelial cells. (Figures 2D and 2E) We identified 2509 arterial ECs, 7357 capillary ECs, 2507 venous ECs, and 2233 lymphatic ECs. The distribution of ECs among the subtypes was statistically different between diets and tissues with the largest difference being observed between the NC Mes and NC Epi ECs (**Supplemental Figure 2A**). Of note, this analysis excludes lymph node ECs which are not present in the adipose endothelium. Lymph node ECs were identified based on their presence only in the mesentery and the expression of several previously established markers of lymph node ECs (**Supplemental Figure 2B and** 2C).^15^

**Figure 2:**
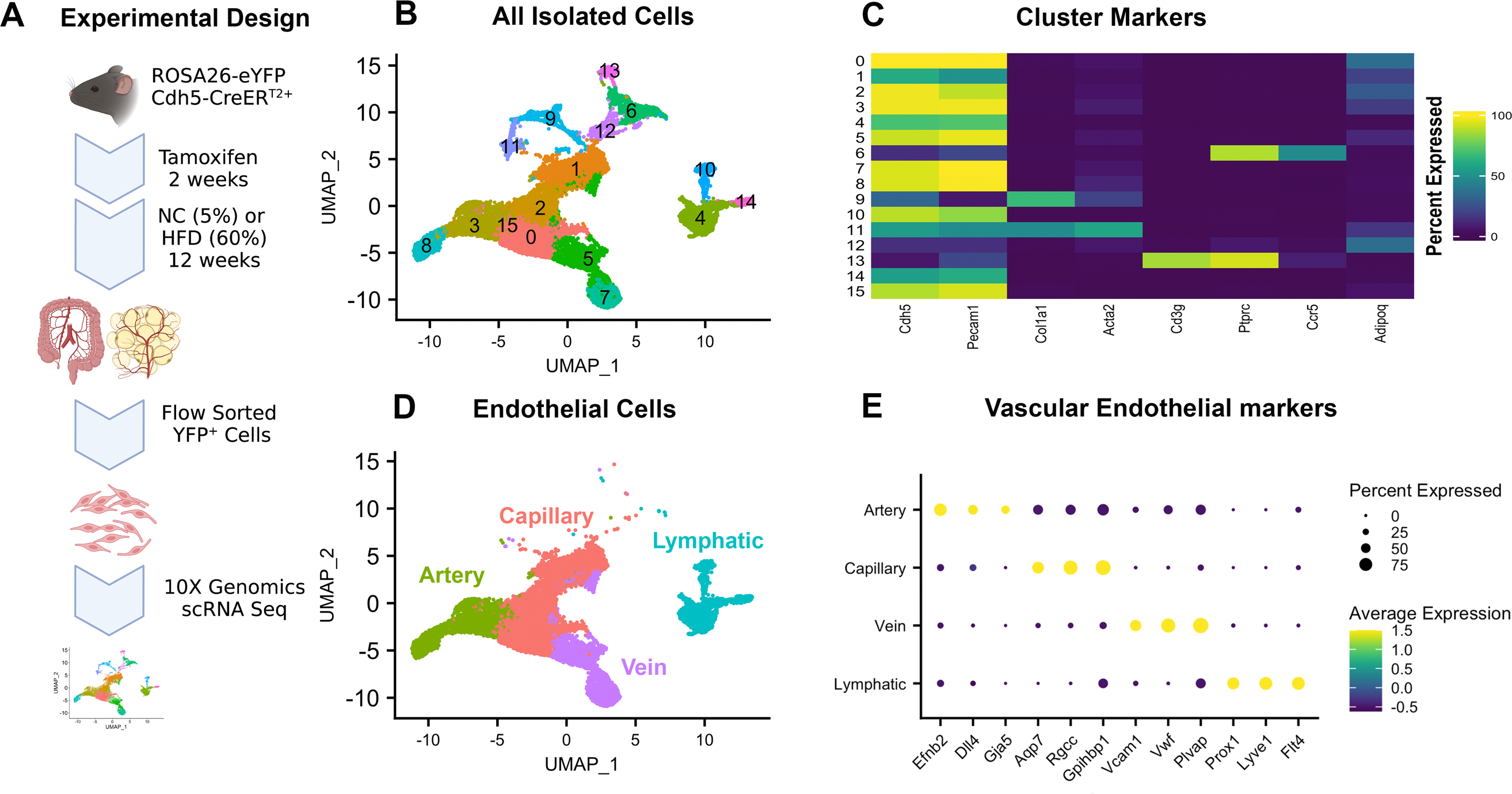
scRNAseq on fluorescently tagged endothelium from adipose and mesenteric vasculature. Experimental design (A) showing the workflow for the isolation of ECs utilizing an inducible YFP reporter mouse under the control of the Cdh5-Cre promoter. Cells were sequenced using 10X genomics. All isolated and sequenced ECs (B) from mesenteric and adipose vasculature are shown with a UMAP consisting of 14 distinct clusters. (C) Contaminating cells (i.e. fibroblasts, smooth muscle cells, immune cells, and adipocytes) not expressing either Cdh5 or Pecam1 were removed from the dataset. (D) UMAP showing only vascular endothelial cells broken into distinct clusters (Artery, Capillary, Vein, Lymphatic) based on vascular markers (E) for each vessel type. scRNAseq on YFP+ ECs from adipose and the mesenteric vasculature cluster together based on vessel type. These data represent two independent scRNAseq experiments (N=2) merged together in Seurat using SCTransform.

To acquire a broad understanding of how a HFD alters the endothelial transcriptome at a single cell resolution, subsets of ECs were isolated, reanalyzed, and plotted using UMAP projections (Figure 3A-D). Again, we removed lymph node ECs from all analysis and only considered lymphatic vessel ECs (**Supplemental Figure 3**). In each vascular subset, the distribution of ECs between diet and tissue was quantified and statistically analyzed using chi-squared tests to compare the magnitude of the differences (Figure 3E-H). In our data, we found Mes and Epi ECs to become more homogenous in their distribution of ECs when exposed to HFD. This difference was most pronounced in arterial ECs (Figure 3H). We additionally used the FindMarkers() function to find differentially expressed genes (DEG; p-value < 0.05 and log2 fold change > 1 or < −1) between Mes and Epi ECs in NC and HFD mice (Figure 3I-L). We found ECs in these two tissues to have fewer DEGs in HFD mice than in NC mice further supporting the convergence of gene expression profiles in endothelium of obese mice. This difference is again most striking in arterial ECs, where in NC mice, ∼7% of genes are differentially expressed between Mes and Epi ECs, but this is reduced to ∼0.5% of genes in HFD fed mice.

**Figure 3:**
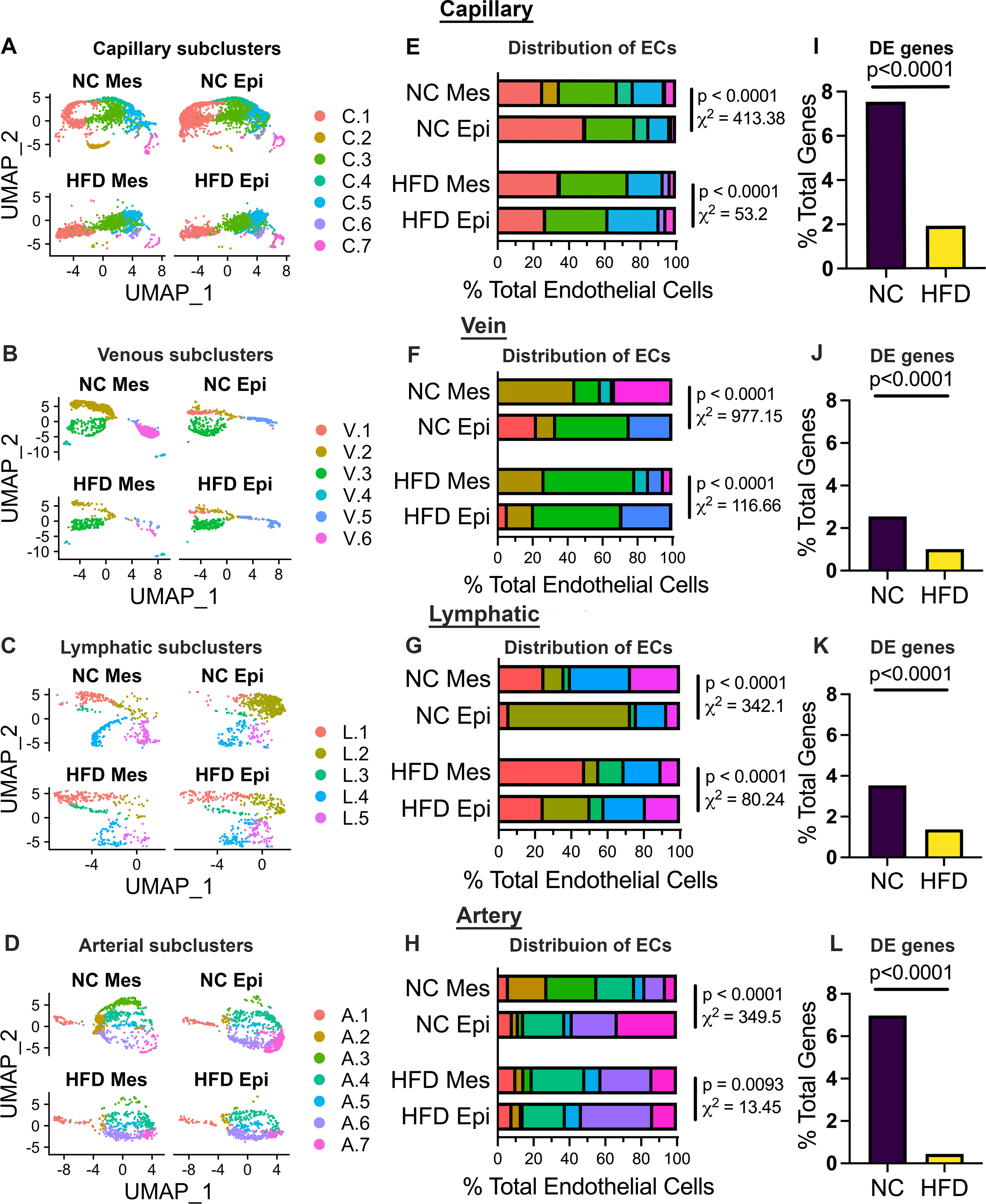
HFD reduces tissue specific EC heterogeneity. Data represented are from both mesenteric (Mes) and epididymal fat (Epi) from mice fed either a normal chow (NC) and high fat diet (HFD). Cells were subsetted and re-clustered into UMAPS based on their location in the vascular tree (A-D). Distributions of these cells for each vascular subtype was quantified and graphed (E-H). Differentially expressed genes between NC and HFD in each vascular subset (Mes and Epi ECs combined) is shown as a percentage of total genes expressed (I-L). The arterial clusters had the biggest decrease in differentially expressed genes (L) from NC to HFD. Chi-squared analysis was used for statistical analysis of cellular distribution.

### Tissue and diet specific mitochondrial gene expression in ECs

The striking convergence seen in arterial ECs combined with the well-established role of arterial dysfunction in cardiovascular disease led to our further exploration of the gene patterns that are changing in response to HFD. Using pathway analysis to identify gene ontology (GO) terms of DEGs in NC ECs but not in HFD ECs, we found several pathways associated with mitochondrial function. (Figure 4A-B **and Supplemental Figure 4**). Specifically, pathways related to mitochondrial electron transport chain were the top pathways most differentially expressed in NC Epi ECs (Figure 4A). In total, 55 of the DEGs encode proteins involved in oxidative phosphorylation, 48 of which were higher in Epi artery ECs compared to Mes artery ECs (Figure 4D). These genes were more lowly expressed in HFD Epi artery ECs showing a similar expression profile to that of HFD Mes artery ECs.

**Figure 4:**
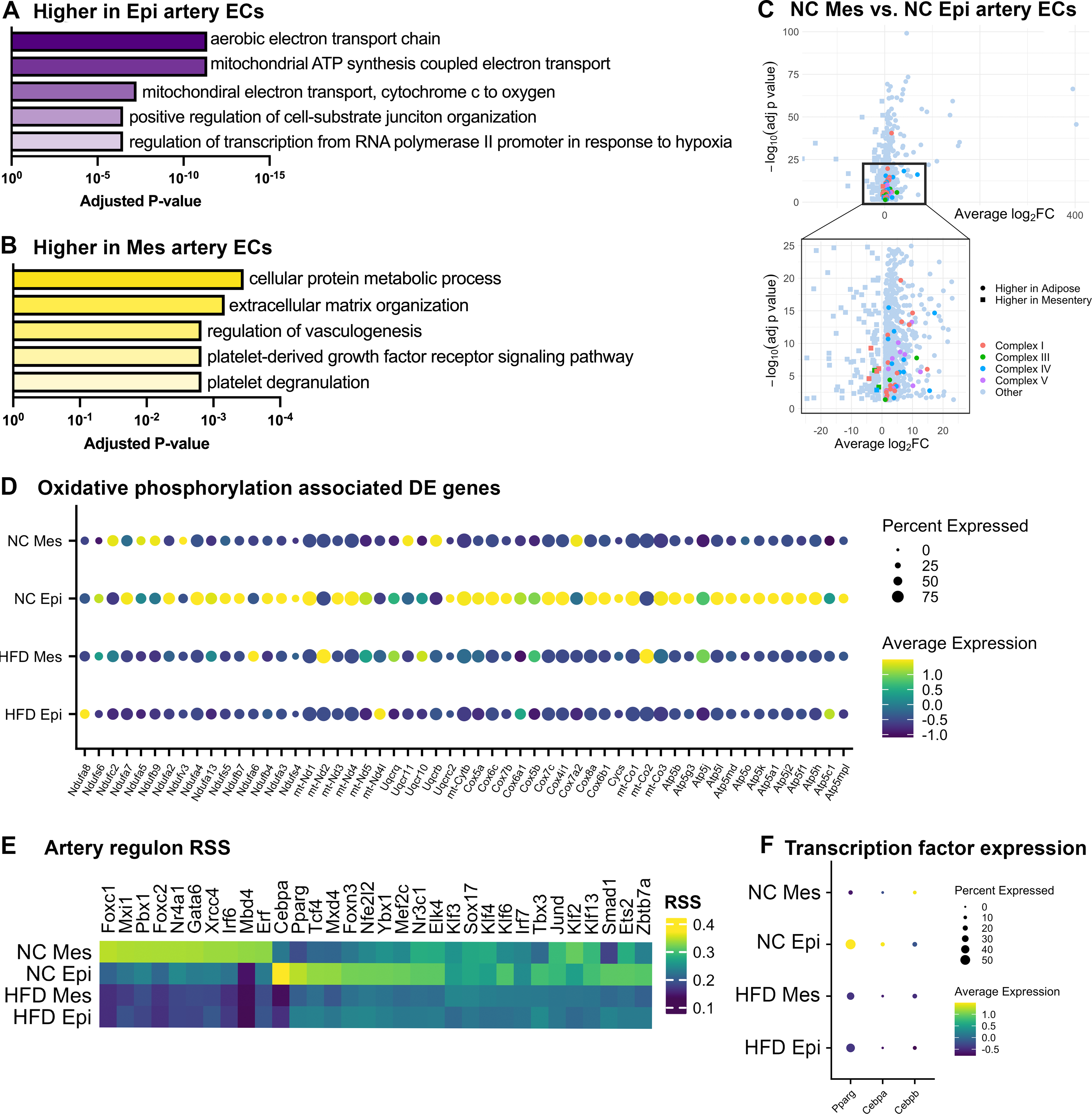
Arterial adipose ECs lose unique metabolic transcriptomic signature during HFD. (A) GO term analysis show increased pathway enrichment for aerobic respiration in epididymal fat pad arteries (Epi) compared to mesenteric arteries (Mes). Mes arteries are metabolically equipped differently than Epi arteries. (B) The majority of the increased pathway enrichment in Mes compared to Epi arteries involves protein metabolism and cell matrix organization. Volcano plot in (C) shows an increased in expression of oxidative phosphorylation related genes in NC Epi arteries when compared to NC Mes arteries. Dotplot (D) displaying several genes associated with oxidative phosphorylation and how the expression (color) and percent of cells (dot size) expressing these genes varies between diet and tissue. (E) SCENIC analysis performed on each sample shows the regulatory networks between tissues and diets converging with HFD. Regulon specificity scores (RSS) have the greatest differences in NC between Mes and Epi. HFD Mes and Epi RSS become more similar. (F) The expression of Pparg and Cebpa are also highest in HFD Epi artery ECs while Cebpb, a related transcription factor, does not follow this pattern.

To better understand what transcriptional pathways dictate these transcriptomic differences, we utilized the SCENIC pipeline.^13^ This analysis infers transcription factor (termed ‘regulon’) activity based on gene expression profiles of each cell. We then calculated a regulon specificity score (RSS) for each group (Figure 4E **and Supplemental Figure 5)** and show the top ten regulons in each group. In alignment with our previous findings, we found NC Mes ECs and NC Epi ECs to have the most distinct regulon activities. Transcription factors with high RSS in either Mes or Epi are decreased in HFD from both tissues. RSS of ECs is more similar between Mes and Epi in HFD than in NC. In arterial ECs, we found C/EBPα (Cebpa) and PPARγ (Pparg) to have the highest RSS in NC Epi ECs. Additionally, expression of Pparg and Cebpa are highest in NC Epi ECs relative to the other groups (Figure 4F). PPARγ and C/EBPα are both known to regulate oxidative phosphorylation and lipid metabolism.^16–18^ Of note, C/EBPβ (Cebpb), which also regulates PPARg and Cebpa in adipocytes^19^, was not identified as having a high RSS nor was it highly expressed in HFD Epi ECs (Figure 4F).

### Human adipose ECs reflect similar diet induces changes as mouse Epi ECs

We next sought to understand the translational relevance of these findings, by utilizing a publicly available snRNA-seq dataset form human visceral adipose tissue containing vascular cells (Figure 5A).^12^ The arterial ECs originally identified in this data set also express the same identifying markers as arterial ECs in our mouse dataset (Figure 4B). We performed SCENIC analysis and calculated RSS of patients grouped based on body mass index (BMI). We found the RSS of PPARγ and C/EBPα to be highest in the group with the lowest BMI, findings that are in agreement with the data from our mouse model (Figure 4C). The regulons with the highest RSS scores of each group are presented in **Supplemental Figure 6**. Additionally, we generated a composite score of genes encoding mitochondrial oxidative phosphorylation proteins using the AddModuleScore() function. The genes used to create this composite score can be found in **Supplementary Table 2**. The composite score (MTscore), PPARG expression, and CEBPA expression all decrease with increasing BMI (Figure 5D).

**Figure 5:**
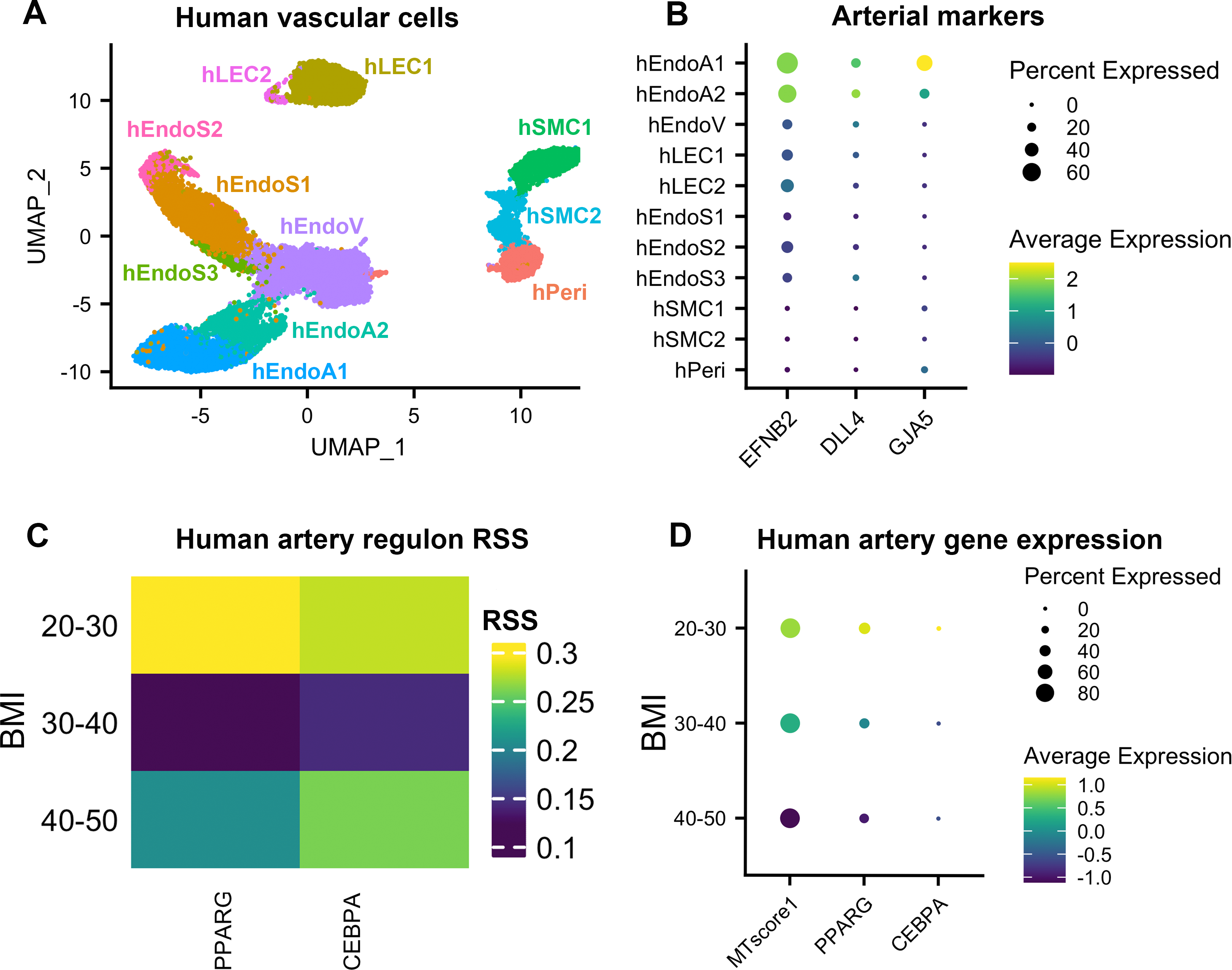
ECs from obese patients show similar transcriptional phenotypes to HFD mice. Human data used here is adapted from Emont et al.^12^ (A) UMAP plot of all sequenced vascular cells from human adipose tissue, samples were taken from both lean and obese patients. (B) Arterial clusters (hEndoA1, hEndoA2) originally defined by the original authors were confirmed using markers EFNB2, DLL4, GJA5. Arterial clusters were then subsetted for further transcriptomic analysis in C-D. (C) Regulon specificity scores (RSS) for top transcription factors found in mouse dataset. (D) Dot plot of key transcription factors PPARG and CEBPA showing decreased expression with increase in BMI, similar to their RSS. MTscore1 is a module score of mitochondrial oxidative phosphorylation genes. A higher MTscore1 indicates higher expression of oxidative phosphorylation genes while a decreased MTscore1 reflects a decrease in oxidative phosphorylation genes. Like PPARG and CEBPA, MTscore1 decreases with an increase in BMI.

### Mitochondrial content confirmed with intact arterial staining

To confirm these findings on the protein level, Mes and Epi arteries were prepared *en face* and immunostained for cytochrome c, a commonly used marker for mitochondria. Cytochrome c is differentially expressed between Mes and Epi arterial ECs in NC fed mice (Figure 4D; Cycs). Arteries from NC Epi fat pads had a higher abundance of cytochrome c staining than all other arteries shown (Figure 6). HFD reduces this increase in cytochrome c, showing a uniform staining pattern between HFD Mes and Epi arteries. Immunofluorescence images were quantified (Figure 6B), using fluorescence intensity.

**Figure 6:**
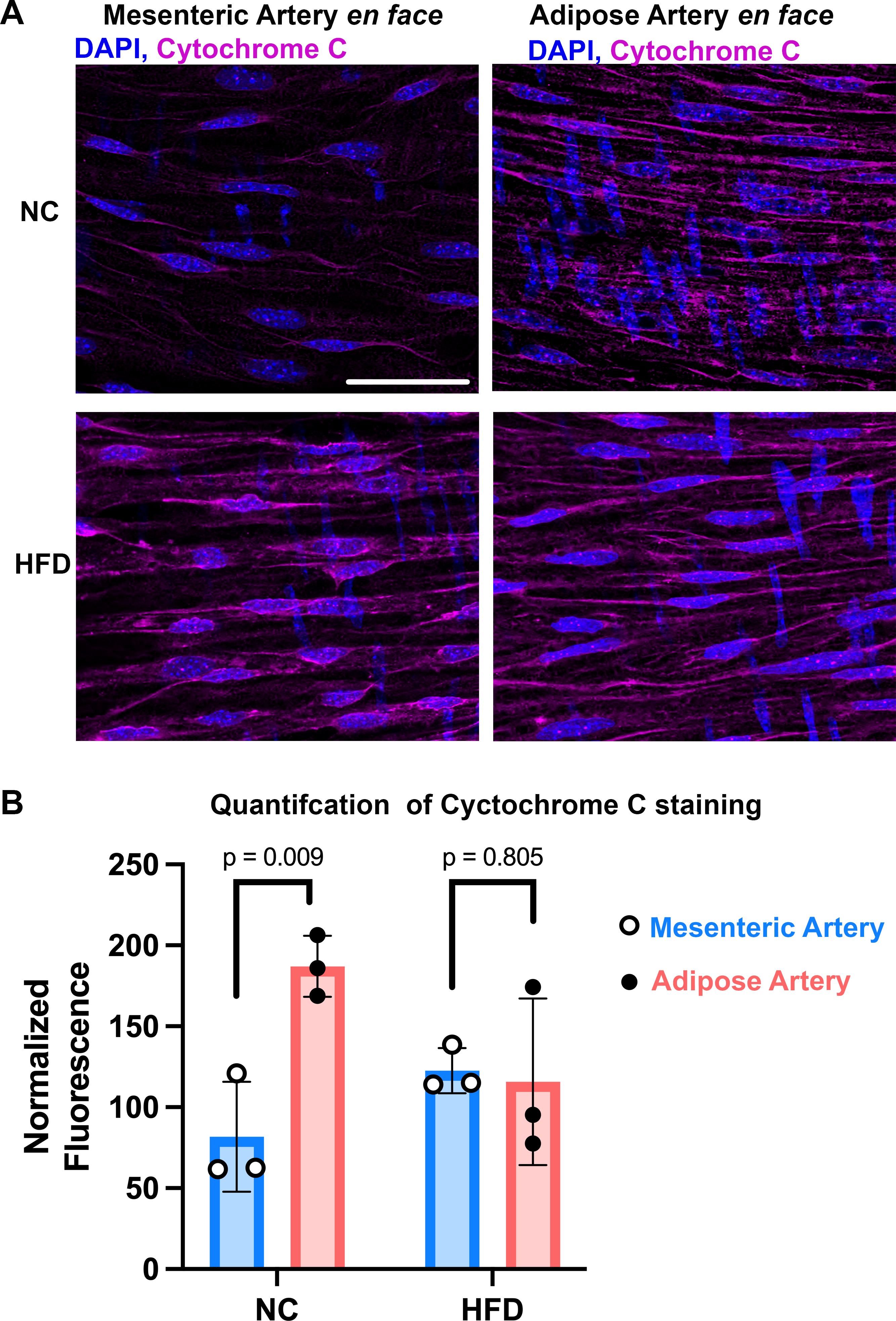
HFD reduced mitochondrial abundance in adipose arteries. *En face* view of both mesenteric and adipose arteries from normal chow (NC) and high fat diet (HFD) fed mice. Nuclei are marked using DAPI (blue) and cytochrome C (mangenta) is used to visualize the mitochondrial network of intact arteries. Fluorescence intensity is quantified in (B) using three images per artery. Statics shown are a two-way ANOVA with a Holm-Sidak post hoc test, N=3 mice. Scale bare = 50 μm for all images.

## Discussion

ECs are a highly heterogenous cell type specialized to suit the function of their resident tissue.^20–22^ Our findings add to this body of knowledge by investigating how EC heterogeneity is impacted by obesity. Specifically, we found adipose ECs to have higher expression of genes involved in mitochondrial respiration and higher mitochondrial content compared to mesenteric ECs in NC fed mice. Furthermore, obesity reduces mitochondrial respiratory gene expression in the adipose endothelium in both mice and humans.

In our mouse model, we examined ECs from the mesentery and epididymal adipose tissue. The mesenteric arteries are commonly used as a model vascular bed due to the vessels contribution to peripheral resistance.^23, 24^ The adipose vasculature regulates tissue blood flow and has recently been implicated in regulating adipocyte metabolism.^25^ We investigated the expression of genes involved in mitochondrial oxidative phosphorylation in isolated ECs with bulk RNA-seq and found Epi ECs to have higher expression of respiratory genes regardless of diet. Bulk RNA-seq, however, is limited to revealing the average expression across a rather heterogenous population of ECs masking potentially meaningful changes in arterial, capillary, venous, or lymphatic ECs. Since ECs are heterogenous and their function is dependent on their location in the vascular tree as well as tissue type (e.g. capillary ECs controlling solute/nutrient transport and arterial ECs controlling tissue perfusion), we hypothesized that ECs would be differentially impacted by an obesogenic diet. To test this hypothesis, we performed scRNA-seq on ECs isolated from mesenteric and adipose tissues. We first investigated the transcriptional heterogeneity of ECs in these two tissues through two methods. First, we subset and remapped each cluster of ECs to quantify the distribution of ECs between clusters. We were able to interpret the distribution of ECs across clusters as a readout of EC similarity due to the nature of UMAP analysis clustering cells of similar transcriptomic signatures in close proximity. The differences between cluster distribution were quantified using the chi-squared test to compare the magnitude heterogeneity. We found that regardless of EC subtype, the distribution of ECs in various clusters from Mes and Epi become more similar with HFD. This is most striking in the arterial endothelium. As a second method, we compared the percentage of genes that were differentially expressed between tissues and found HFD decreased the percentage of DEGs in each subset of ECs. The largest decrease in DEGs was seen in arterial ECs.

We then investigated what pathways are most affected in the arterial endothelium in response to obesity. Genes that were differentially expressed in NC but not HFD were used for gene ontology (GO) analysis revealing respiratory associated genes. Specifically, genes associated with oxidative phosphorylation and electron transport chain pathways were more highly expressed in Epi than Mes. Interestingly, an obesogenic diet diminishes this difference where HFD Mes and Epi ECs have reduced expression of respiratory pathway associated genes. These data show an apparent contradiction to the results of our bulk RNA-seq data where Epi ECs had higher oxidative phosphorylation gene expression regardless of diet. This demonstrates the limitation of bulk RNA-seq while exemplifying the benefits of scRNAseq. Our scRNAseq data revealed intratissue EC heterogeneity that was previously masked by bulk RNA-seq.

To confirm our mRNA-based findings, we isolated both mesenteric and adipose arteries and prepared them *en face* to quantify endothelial mitochondrial content. To our knowledge, this is the first time an adipose artery has been prepared and viewed *en face*, as well as the first time cytochrome c staining has been performed in an intact endothelium. Our protein staining confirms our findings by showing adipose arteries have higher abundance of cytochrome c than mesenteric arteries, and HFD reduces cytochrome C content in adipose arteries such that they become similar to mesenteric arteries.

These findings are interesting to consider in light of recent debate on the dependence of endothelial cells on glycolysis or mitochondrial respiration. Our results suggest a greater dependence of adipose artery ECs on oxidative phosphorylation than mesenteric artery ECs in physiologic conditions. While we have not investigated the mechanism(s) that result in this heterogeneity, Monelli and colleagues have proposed a model by which free fatty acid availability from local adipocyte lipolysis can regulate adipose EC metabolism.^25^ It may be adipose artery ECs are simply responding to the local availability of substrate for oxidative phosphorylation and reduce the expression of these genes when lipolysis decreases in HFD.^26, 27^ When considering adipocyte-generated factors as a potential mechanism, it is quite surprising that arterial ECs display the largest degrees of convergence. One might expect the ECs in closest proximity with adipocytes (i.e. capillary ECs), immediately downstream of the adipocytes (i.e. venous ECs), or in proximity to adipose interstitial fluid (i.e. lymphatic ECs) to experience the highest concentrations adipokines and therefore be most affected. It therefore may be changes in blood flow or signaling via smooth muscle cells which drive the changes seen in arterial ECs.

We investigated which transcription factors are involved in this convergence. SCENIC analysis of these data revealed C/EBPα and PPARγ, both involved in lipid metabolism,^16, 17^ had the highest RSS and gene expression in NC Epi ECs. In adipocytes, both of these transcription factors are regulated by C/EBPβ.^19^ However, we did not observe increased expression or high RSS of C/EBPβ in NC Epi ECs suggesting C/EBPα and PPARγ expression is induced through C/EBPβ independent mechanisms in ECs.

PPARγ agonists are used to improve insulin sensitivity in patients with type two diabetes and improve endothelial function.^28^ PPARγ agonists have been shown to increase mitochondrial content of adipocytes^18^ and increase fatty acid import genes in the endothelium.^29^ However, to our knowledge, whether these agonists increase arterial EC mitochondrial content has not been investigated. Our findings suggest PPARγ agonism may change EC function through increasing mitochondrial content.

To examine the translatability of our findings, we used a published dataset^12^ to investigate these pathways in obese human visceral adipose ECs. We created a module score of oxidative phosphorylation genes and found this score decreased with increasing BMI. Similarly, both CEBPA and PPARG expression and RSS decreased with obesity suggesting these findings also hold true in human obesity.

Our findings are limited by the exclusion of female mice. However, the human data contain both male and female subjects with an over representation of females. While these data are under-powered to analyze sex differences, the fact that the findings of our mouse model were recapitulated in this human data set suggests obesity reduces adipose artery EC mitochondrial content in both males and females. Further work is necessary to thoroughly investigate sex differences in adipose EC mitochondrial content.

Arterial EC dysfunction is prevalent in diet-induced obesity and associated with cardiovascular disease progression.^1–5^ Here we show obesity reduces mitochondrial content in adipose artery ECs resulting in a loss of tissue specific EC heterogeneity. Our findings pave the way for future work to further investigate vascular (mal)adaptations to obesity.

## Non-standard Abbreviations and Acronyms

EC: Endothelial Cell
NC: Normal Chow
HFD: Hight Fat Diet
Mes: Mesenteric
Epi: Epididymal fat pad
RSS: Regulon Specificity Score
RNA-seq: RNA Sequencing
scRNA-seq: Single Cell RNA Sequencing
snRNA-seq: Single Nuclei RNA Sequencing
SCENIC: Single-Cell Regulatory Network Inference and Clustering

## Acknowledgments

We thank School of Medicine Genome Analysis and Technology Core, RRID:SCR_018883 and the University of Virginia Flow Cytometry Core, RRID: SCR_017829 for their technical expertise.

## Sources of Funding

University of Virginia Basic and Translational Cardiovascular Research Training Grant T32 007284 (L.S.D. and M.A.L.); NIH HL137112 (B.E.I.); University of Virginia LaunchPad (B.E.I.); Lipedema Foundation (B.E.I.); AHA Predoctoral Fellowship #915176 (M.A.L.)

## Disclosures

None.

## Supplemental Material

Figures S1-S6

Tables S1-S2

## Notes

https://www.ncbi.nlm.nih.gov/geo/query/acc.cgi?acc=GSE235192

### Competing Interest Statement

The authors have declared no competing interest.

## References

1. van der Heijden DJ, van Leeuwen MAH, Janssens GN, Lenzen MJ, van de Ven PM, Eringa EC and van Royen N. Body Mass Index Is Associated With Microvascular Endothelial Dysfunction in Patients With Treated Metabolic Risk Factors and Suspected Coronary Artery Disease. J Am Heart Assoc. 2017;6.

2. Gimbrone MA, Jr. and Garcia-Cardena G. Endothelial Cell Dysfunction and the Pathobiology of Atherosclerosis. Circ Res. 2016;118:620–36.

3. Kwasniewska M, Kozinska J, Dziankowska-Zaborszczyk E, Kostka T, Jegier A, Rebowska E, Orczykowska M, Leszczynska J and Drygas W. The impact of long-term changes in metabolic status on cardiovascular biomarkers and microvascular endothelial function in middle-aged men: a 25-year prospective study. Diabetol Metab Syndr. 2015;7:81.

4. Kavurma MM, Bursill C, Stanley CP, Passam F, Cartland SP, Patel S, Loa J, Figtree GA, Golledge J, Aitken S and Robinson DA. Endothelial cell dysfunction: Implications for the pathogenesis of peripheral artery disease. Front Cardiovasc Med. 2022;9:1054576.

5. Brainin P, Frestad D and Prescott E. The prognostic value of coronary endothelial and microvascular dysfunction in subjects with normal or non-obstructive coronary artery disease: A systematic review and meta-analysis. Int J Cardiol. 2018;254:1–9.

6. Krutzfeldt A, Spahr R, Mertens S, Siegmund B and Piper HM. Metabolism of exogenous substrates by coronary endothelial cells in culture. J Mol Cell Cardiol. 1990;22:1393–404.

7. Dobrina A and Rossi F. Metabolic properties of freshly isolated bovine endothelial cells. Biochim Biophys Acta. 1983;762:295–301.

8. Quintero M, Colombo SL, Godfrey A and Moncada S. Mitochondria as signaling organelles in the vascular endothelium. Proc Natl Acad Sci U S A. 2006;103:5379–84.

9. Ibrahim A, Yucel N, Kim B and Arany Z. Local Mitochondrial ATP Production Regulates Endothelial Fatty Acid Uptake and Transport. Cell Metab. 2020;32:309–319 e7.

10. Schiffmann LM, Werthenbach JP, Heintges-Kleinhofer F, Seeger JM, Fritsch M, Gunther SD, Willenborg S, Brodesser S, Lucas C, Jungst C, Albert MC, Schorn F, Witt A, Moraes CT, Bruns CJ, Pasparakis M, Kronke M, Eming SA, Coutelle O and Kashkar H. Mitochondrial respiration controls neoangiogenesis during wound healing and tumour growth. Nat Commun. 2020;11:3653.

11. Wilson C, Lee MD, Buckley C, Zhang X and McCarron JG. Mitochondrial ATP Production is Required for Endothelial Cell Control of Vascular Tone. Function (Oxf). 2023;4:zqac063.

12. Emont MP, Jacobs C, Essene AL, Pant D, Tenen D, Colleluori G, Di Vincenzo A, Jorgensen AM, Dashti H, Stefek A, McGonagle E, Strobel S, Laber S, Agrawal S, Westcott GP, Kar A, Veregge ML, Gulko A, Srinivasan H, Kramer Z, De Filippis E, Merkel E, Ducie J, Boyd CG, Gourash W, Courcoulas A, Lin SJ, Lee BT, Morris D, Tobias A, Khera AV, Claussnitzer M, Pers TH, Giordano A, Ashenberg O, Regev A, Tsai LT and Rosen ED. A single-cell atlas of human and mouse white adipose tissue. Nature. 2022;603:926–933.

13. Aibar S, Gonzalez-Blas CB, Moerman T, Huynh-Thu VA, Imrichova H, Hulselmans G, Rambow F, Marine JC, Geurts P, Aerts J, van den Oord J, Atak ZK, Wouters J and Aerts S. SCENIC: single-cell regulatory network inference and clustering. Nat Methods. 2017;14:1083–1086.

14. Suo S, Zhu Q, Saadatpour A, Fei L, Guo G and Yuan GC. Revealing the Critical Regulators of Cell Identity in the Mouse Cell Atlas. Cell Rep. 2018;25:1436–1445 e3.

15. Jalkanen S and Salmi M. Lymphatic endothelial cells of the lymph node. Nat Rev Immunol. 2020;20:566–578.

16. Briot A, Decaunes P, Volat F, Belles C, Coupaye M, Ledoux S and Bouloumie A. Senescence Alters PPARgamma (Peroxisome Proliferator-Activated Receptor Gamma)-Dependent Fatty Acid Handling in Human Adipose Tissue Microvascular Endothelial Cells and Favors Inflammation. Arterioscler Thromb Vasc Biol. 2018;38:1134–1146.

17. Zaini MA, Muller C, de Jong TV, Ackermann T, Hartleben G, Kortman G, Guhrs KH, Fusetti F, Kramer OH, Guryev V and Calkhoven CF. A p300 and SIRT1 Regulated Acetylation Switch of C/EBPalpha Controls Mitochondrial Function. Cell Rep. 2018;22:497–511.

18. Bogacka I, Xie H, Bray GA and Smith SR. Pioglitazone induces mitochondrial biogenesis in human subcutaneous adipose tissue in vivo. Diabetes. 2005;54:1392–9.

19. Ghaben AL and Scherer PE. Adipogenesis and metabolic health. Nat Rev Mol Cell Biol. 2019;20:242–258.

20. Kalucka J, de Rooij L, Goveia J, Rohlenova K, Dumas SJ, Meta E, Conchinha NV, Taverna F, Teuwen LA, Veys K, Garcia-Caballero M, Khan S, Geldhof V, Sokol L, Chen R, Treps L, Borri M, de Zeeuw P, Dubois C, Karakach TK, Falkenberg KD, Parys M, Yin X, Vinckier S, Du Y, Fenton RA, Schoonjans L, Dewerchin M, Eelen G, Thienpont B, Lin L, Bolund L, Li X, Luo Y and Carmeliet P. Single-Cell Transcriptome Atlas of Murine Endothelial Cells. Cell. 2020;180:764–779 e20.

21. Aird WC. Phenotypic heterogeneity of the endothelium: II. Representative vascular beds. Circ Res. 2007;100:174–90.

22. Aird WC. Phenotypic heterogeneity of the endothelium: I. Structure, function, and mechanisms. Circ Res. 2007;100:158–73.

23. Wenceslau CF, McCarthy CG, Earley S, England SK, Filosa JA, Goulopoulou S, Gutterman DD, Isakson BE, Kanagy NL, Martinez-Lemus LA, Sonkusare SK, Thakore P, Trask AJ, Watts SW and Webb RC. Guidelines for the measurement of vascular function and structure in isolated arteries and veins. Am J Physiol Heart Circ Physiol. 2021;321:H77–H111.

24. Christensen KL and Mulvany MJ. Mesenteric arcade arteries contribute substantially to vascular resistance in conscious rats. J Vasc Res. 1993;30:73–9.

25. Monelli E, Villacampa P, Zabala-Letona A, Martinez-Romero A, Llena J, Beiroa D, Gouveia L, Chivite I, Zagmutt S, Gama-Perez P, Osorio-Conles O, Muixi L, Martinez-Gonzalez A, Castillo SD, Martin-Martin N, Castel P, Valcarcel-Jimenez L, Garcia-Gonzalez I, Villena JA, Fernandez-Ruiz S, Serra D, Herrero L, Benedito R, Garcia-Roves P, Vidal J, Cohen P, Nogueiras R, Claret M, Carracedo A and Graupera M. Angiocrine polyamine production regulates adiposity. Nat Metab. 2022;4:327–343.

26. Bougneres P, Stunff CL, Pecqueur C, Pinglier E, Adnot P and Ricquier D. In vivo resistance of lipolysis to epinephrine. A new feature of childhood onset obesity. J Clin Invest. 1997;99:2568–73.

27. Kasher-Meron M, Youn DY, Zong H and Pessin JE. Lipolysis defect in white adipose tissue and rapid weight regain. Am J Physiol Endocrinol Metab. 2019;317:E185–E193.

28. Nesti L, Trico D, Mengozzi A and Natali A. Rethinking pioglitazone as a cardioprotective agent: a new perspective on an overlooked drug. Cardiovasc Diabetol. 2021;20:109.

29. Goto K, Iso T, Hanaoka H, Yamaguchi A, Suga T, Hattori A, Irie Y, Shinagawa Y, Matsui H, Syamsunarno MR, Matsui M, Haque A, Arai M, Kunimoto F, Yokoyama T, Endo K, Gonzalez FJ and Kurabayashi M. Peroxisome proliferator-activated receptor-gamma in capillary endothelia promotes fatty acid uptake by heart during long-term fasting. J Am Heart Assoc. 2013;2:e004861.

