## Supplemental Figures for "Loss of endothelial cell heterogeneity in arteries after obesogenic diet"

**SUPPLEMENTARY MATERIAL FOR:**
### Loss of endothelial cell heterogeneity in arteries after obesogenic diet

Luke S. Dunaway, PhD[1,*]
Melissa A. Luse, MS[1,2,*]
Shruthi Nyshadham[1]
Gamze Bulut, PhD[1]
Gabriel F. Alencar, PhD[3]
Nicholas W. Chavkin, PhD[1,4]
Miriam Cortese-Krott[5]
Karen K. Hirschi, PhD[1,4]
Brant E. Isakson, PhD[1,2,#]

[1]Robert M. Berne Cardiovascular Research Center, University of Virginia School of Medicine
[2]Department of Molecular Physiology and Biophysics, University of Virginia School of Medicine
[3]Department of Biochemistry and Molecular Genetics, University of Virginia School of Medicine, Charlottesville, VA, USA
[4]Department of Cell Biology, University of Virginia School of Medicine
[5]Department of Cardiology, Pneumology and Angiology, Heinrich Heine University of Düsseldorf, Düsseldorf Germany

* these authors contributed equally to this work
[#] to whom correspondence should be addressed
University of Virginia School of Medicine
PO Box 801394
Charlottesville, VA 22908
E:
P: 434-924-2093

Supplemental Figures: 6
Supplemental Tables: 2

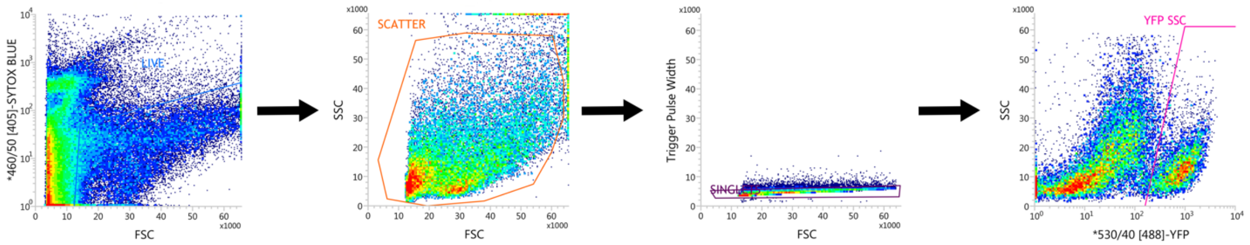


**Supplemental Figure 1: Representative fluorescence activated cell sorting panel**. Cells were sorted on SYTOX blue, forward scatter (FSC), side scatter (SSC), trigger pulse width, and YFP to isolate live, singlet, YFP+ cells.

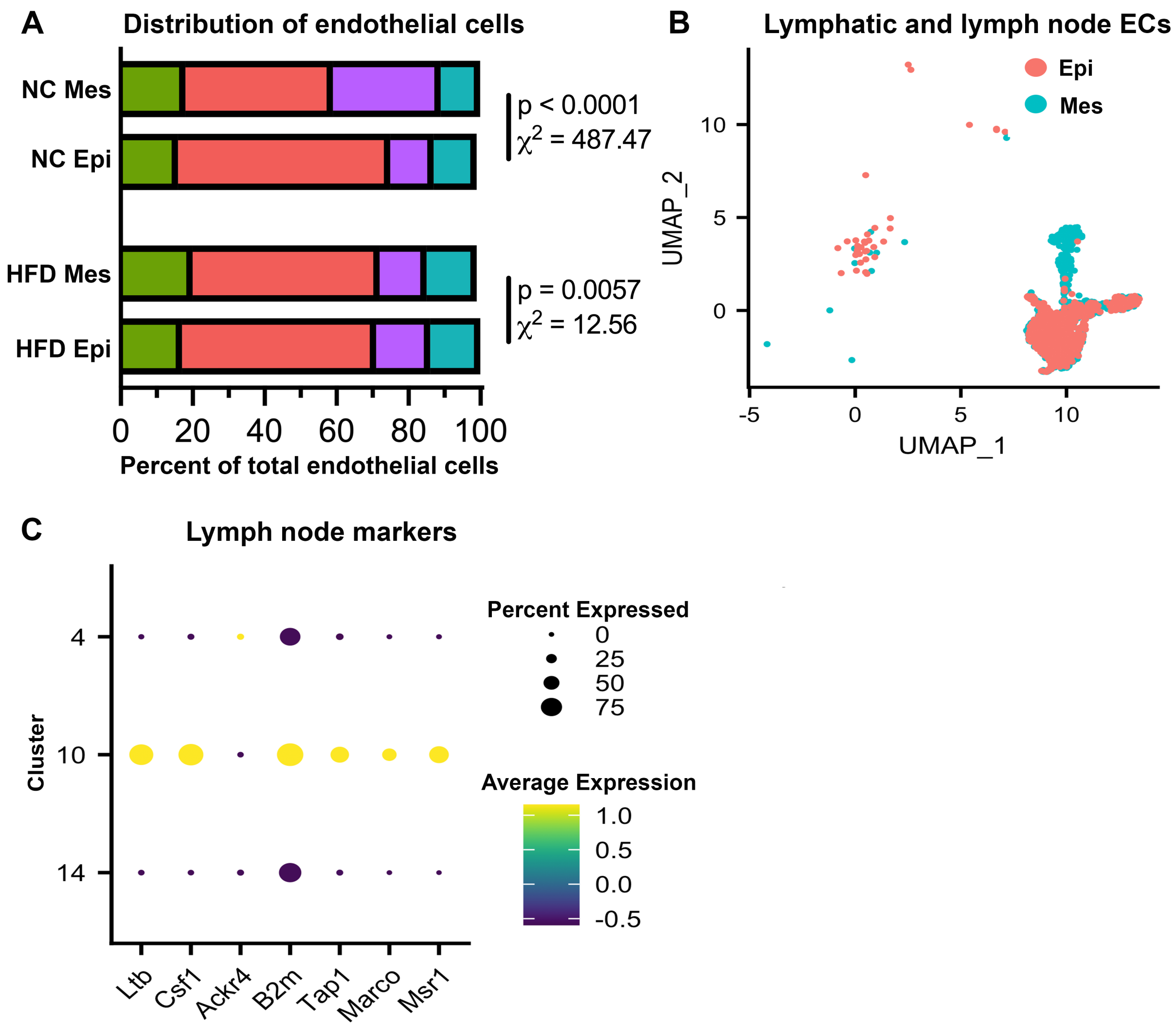


**Supplemental Figure 2: Distribution and exclusion of ECs between diet and tissue.** (A) Bar plot representing the arterial (green), capillary (pink), vein (purple), and lymphatic (blue) breakdown in each tissue and diet. Data are shown as percent of total endothelial cells to account for varying cell number between tissue and diets. Corresponding UMAP can be seen in 2D. (B) UMAP comprising of all lymphatic ECs split between Mes (blue) and Epi (pink). Lymphatic ECs specifically from lymph nodes (C: cluster 10) were characterized by lymph node markers and excluded from this study.

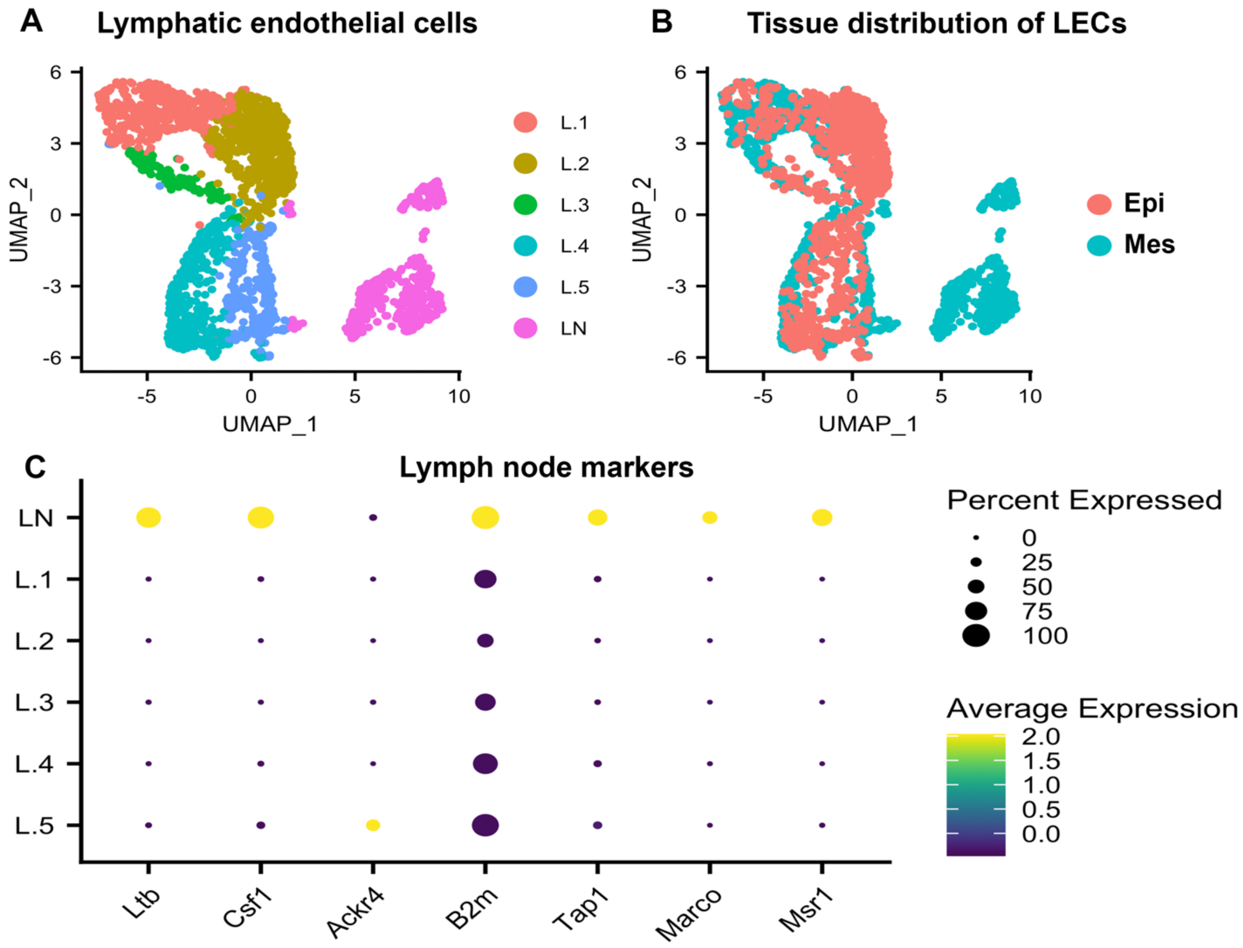


**Supplemental Figure 3: Exclusion of lymph node ECs.** UMAP representing all sequenced lymphatic ECs (A) using markers Prox1, Lyve1, Flt4 (defined in Figure2E). (B) Shows the tissue distribution of lymphatic ECs between adipose (Epi) and Mesenteric (Mes) tissue. (C) Markers used to exclude lymph node ECs from the vascular EC data set.

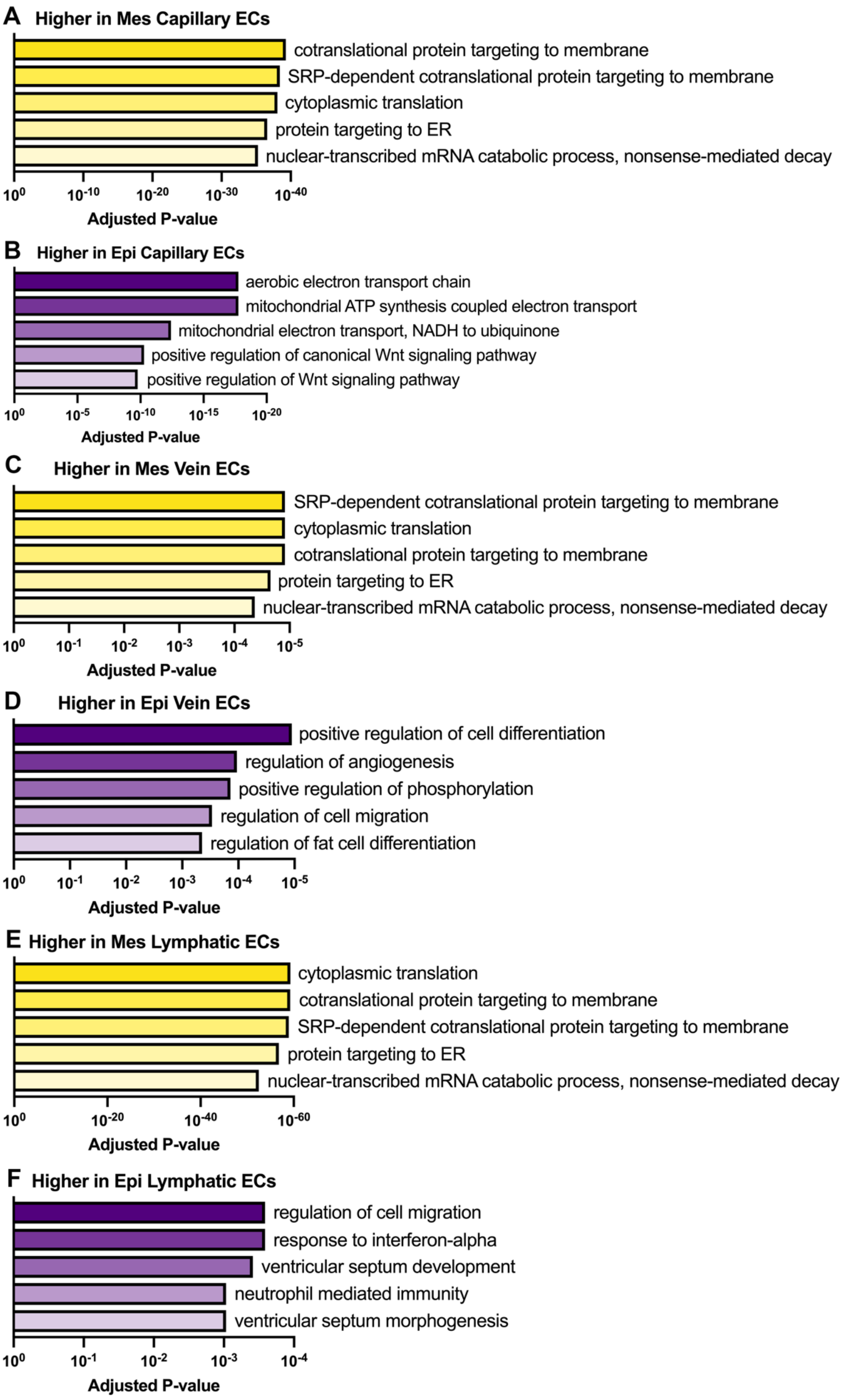
A
Higher in Mes Capillary ECs
cotranslational protein targeting to membrane
SRP-dependent cotranslational protein targeting to membrane
cytoplasmic translation
protein targeting to ER
nuclear-transcribed mRNA catabolic process, nonsense-mediated decay
$10^{0}$ $10^{-10}$ $10^{-20}$ $10^{-30}$ $10^{-40}$
Adjusted P-value
B
Higher in Epi Capillary ECs
aerobic electron transport chain
mitochondrial ATP synthesis coupled electron transport
mitochondrial electron transport, NADH to ubiquinone
positive regulation of canonical Wnt signaling pathway
positive regulation of Wnt signaling pathway
$10^{0}$ $10^{-5}$ $10^{-10}$ $10^{-15}$ $10^{-20}$
Adjusted P-value
C
Higher in Mes Vein ECs
SRP-dependent cotranslational protein targeting to membrane
cytoplasmic translation
cotranslational protein targeting to membrane
protein targeting to ER
nuclear-transcribed mRNA catabolic process, nonsense-mediated decay
$10^{0}$ $10^{-1}$ $10^{-2}$ $10^{-3}$ $10^{-4}$ $10^{-5}$
Adjusted P-value
D
Higher in Epi Vein ECs
positive regulation of cell differentiation
regulation of angiogenesis
positive regulation of phosphorylation
regulation of cell migration
regulation of fat cell differentiation
$10^{0}$ $10^{-1}$ $10^{-2}$ $10^{-3}$ $10^{-4}$ $10^{-5}$
Adjusted P-value
E
Higher in Mes Lymphatic ECs
cytoplasmic translation
cotranslational protein targeting to membrane
SRP-dependent cotranslational protein targeting to membrane
protein targeting to ER
nuclear-transcribed mRNA catabolic process, nonsense-mediated decay
$10^{0}$ $10^{-20}$ $10^{-40}$ $10^{-60}$
Adjusted P-value
F
Higher in Epi Lymphatic ECs
regulation of cell migration
response to interferon-alpha
ventricular septum development
neutrophil mediated immunity
ventricular septum morphogenesis
$10^{0}$ $10^{-1}$ $10^{-2}$ $10^{-3}$ $10^{-4}$
Adjusted P-value

**Supplemental Figure 4: GO terms by vascular bed and tissue.** Go analysis performed on all vascular clusters. GO terms are split as either higher in mesentery (Mes) or higher in adipose (Epi). This analysis was done for capillaries (A-B), Veins (C-D), and Lymphatics (D-E). All p-values are listed on the Y-axis and expressed as adjusted p-value by Mann-Whitney U test.

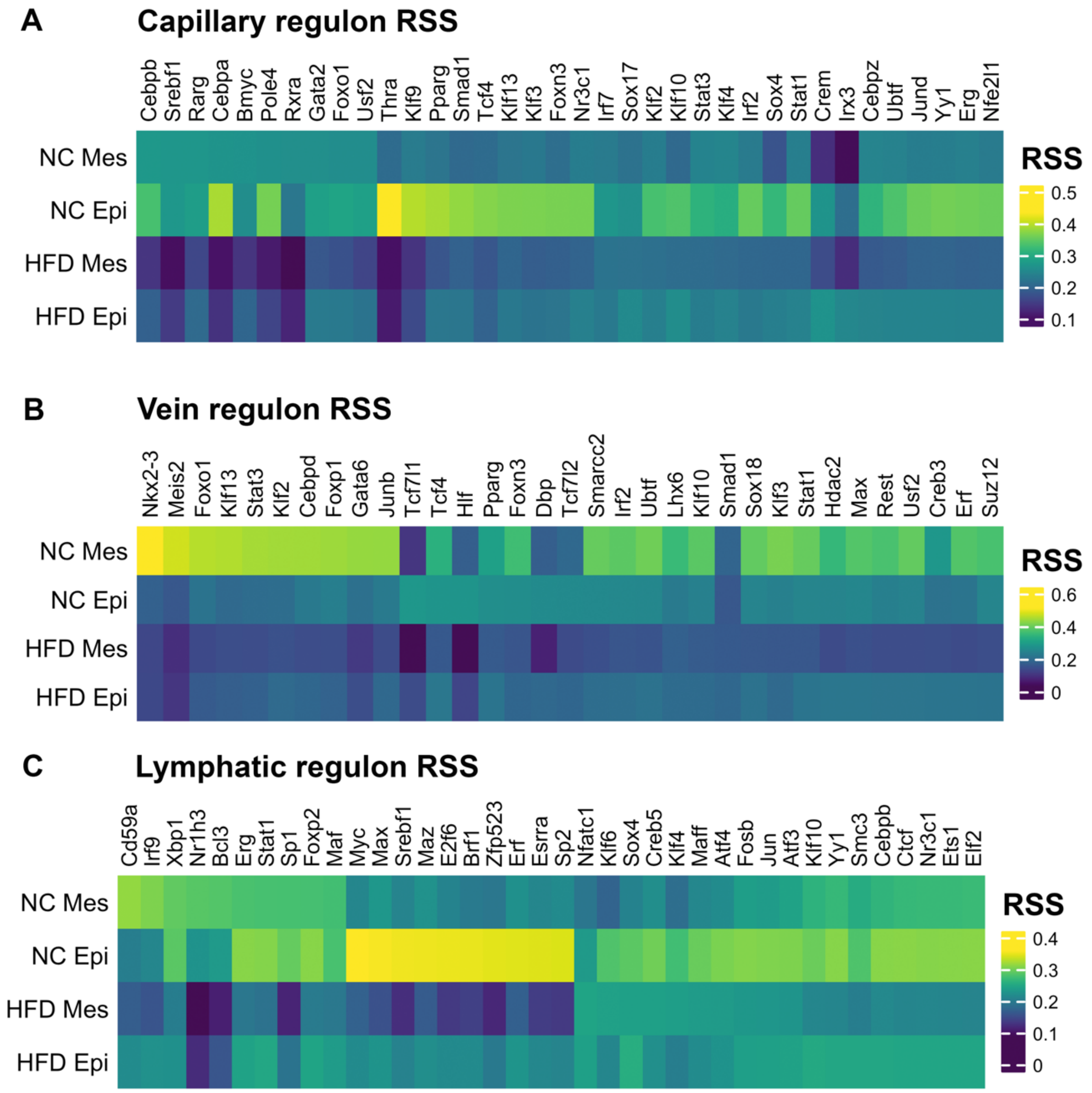


**Supplemental Figure 5: Regulon activity in different vascular beds.** SCENIC analysis was performed on all vascular clusters. Regulon specificity scores (RSS) were assigned to key transcription factors found in the regulon network for each tissue and diet. Capillary regulons (A), vein regulons (B), and lymphatic regulons (C) are shown with yellow as a higher and purple as a lower RSS respectively.

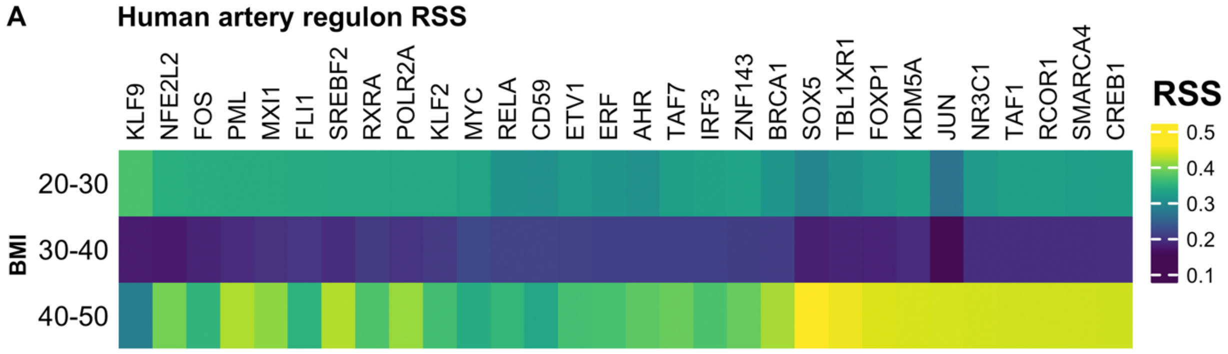


**Supplemental Figure 6: Regulon activity in human arterial ECs.** Regulon network analysis performed on human adipose ECs from patients with various BMIs reflect changes in regulon specificity scores (RSS) as BMI increases.
